# MicroRNA modulation of viral nervous necrosis resistance in European seabass

**DOI:** 10.64898/2026.02.16.706017

**Authors:** Rodríguez-Vázquez Raquel, Robert Mukiibi, Serena Ferraresso, Rafaella Franch, Luca Peruzza, Giulia Dalla Rovere, Jelena Radojicic, Massimiliano Babbucci, Daniela Bertotto, Anna Toffan, Francesco Pascoli, Carolina Peñaloza, Ross D. Houston, Costas S. Tsigenopoulos, Luca Bargelloni, Diego Robledo

## Abstract

MicroRNAs (miRNAs) are key post-transcriptional regulators of antiviral immunity, controlling gene expression by targeting 3’ UTRs of immune-related transcripts. Despite their importance, the role of miRNAs in viral nervous necrosis (VNN) resistance in European seabass (*Dicentrarchus labrax*) is unexplored. Here, we characterized for the first time the brain miRNome of seabass from three VNN-resistance genotypes (susceptible, intermediate, resistant) across two genetically distinct seabass clusters. Differential expression analyses revealed cluster-specific patterns, with susceptible fish consistently showing overexpression of the differently expressed miRNAs (DEmiRNAs) as compared to the resistant fish. Considering the two genetic clusters in the study, miR–199–5p was differentially expressed between the VNN susceptible and resistant fish. This miRNA was found to be less expressed in the resistant individuals. Functional characterization of the miRNA predicted that it binds to two distinct miRNA recognition elements (MREs) within the *ifi27l2a* 3’ UTR. These MREs flank a SNP (Chr3:10,082,380) previously associated with VNN survival. A strong negative correlation (r= -0.840) between miR–199–5p expression and *ifi27l2a* mRNA abundance further supports a post–transcriptional repression mechanism. Together, these results propose a regulatory model in which miR–199–5p modulates *ifi27l2a* expression, contributing to phenotypic variation in VNN resistance and positioning it as a promising biomarker for seabass aquaculture breeding.

## INTRODUCTION

The European seabass (*Dicentrarchus labrax*) is one of the most important marine fish in European and Mediterranean aquaculture, with a significant impact on the regional socio-economic landscape (Vandeputte et al., 2019). Global production has reached approximately 372,000 tonnes in recent years (Saif et al., 2025), with projections estimating an increase to nearly 130 million tonnes of aquatic proteins by 2050 to meet rising demand (Boyd et al., 2022; Costello et al., 2020). Despite this growth, diseases continue to represent a major challenge to the industry, resulting in a significant mortality rate thus causing substantial economic losses (Boyd et al., 2022; Costello et al., 2020). Therefore, enhancing resistance to diseases is a pivotal aspect in the seabass aquaculture industry. Disease challenges in aquaculture are managed via use of antimicrobial compounds, high-quality feeds and vaccines and selective breeding for disease resistance has been proven to be a sustainable and effective approach to enhancing the overall health of fish and reducing economic losses (Buchmann, 2022).

Nervous necrosis virus (NNV) is one of the most virulent pathogens in the aquaculture industry, a single-stranded positive-sense RNA virus which belongs to the *betanodavirus* genus of the family *nodaviridade* (Mori et al., 1992). This highly contagious causative agent is responsible for viral nervous necrosis (VNN), a highly fatal, contagious viral infection that is a key viral disease affecting multiple species of marine and freshwater fishes and accounts for 15% of infectious mortalities in European seabass (Bandín & Souto, 2020; Muniesa et al., 2020). NNV primarily targets the central nervous system tissues (mainly the brain, spinal cord and the retina), where viral replication and multiplication occurs (Chi et al., 2016). Infected tissues are characterized by the presence of atypical abnormalities in numerous vacuolar lesions (Munday et al., 2002). This disease has been responsible for significant mortality rates and considerable economic losses among fish species on a global scale (Moin et al., 2024).

Recently research by Mukiibi et al. 2025 has identified a major quantitative trait loci (QTL) for VNN resistance in Europea seabass on chromosome 3 which confers around 90% of survival against a VNN outbreak. This QTL contains two copies of interferon alpha inducible protein 27-like 2A (*ifi27l2a*) genes and one *ifi27l2,* whose differential expression has been linked to variation in VNN resistance. Importantly, SNPs located within the 3’ UTR of *ifi27l2a* suggest that post–transcriptional regulation, potentially mediated by microRNAs, may play a key role in modulating this antiviral gene (Mukiibi et al., 2025).

Small non-coding RNAs (sncRNAs) including microRNAs (miRNAs) represent an important category of epigenetic regulators of gene expression in cells that consequently contribute to phenotypic variability of different traits. These non-coding RNA molecules with lengths of length of 18-24 nucleotides, have been shown to exert a regulatory influence on gene expression at the post-transcriptional level (Bartel, 2004). They are highly conserved phylogenetically and hundreds of conserved genes encoding miRNAs have been found in various organisms (Ambros et al., 2003). These miRNAs regulate gene expression by binding to miRNA-responsive elements (MREs) in the three prime untranslated regions (3’ UTRs) of target mature mRNAs. They function as gene regulators through base-pairing interactions with their target mRNAs, typically involving a complementary sequence of about 8 nucleotides within the seed region that mediates mRNA degradation or translational repression (Cai et al., 2009; Diener et al., 2024). Furthermore, one mRNA could be the target of multiple miRNAs and one miRNA could interact with multiple mRNAs (Peter, 2010). Involvement of miRNAs in modulating both the innate and adaptive immune systems is well established in mammals (Mehta & Baltimore, 2016), however there is limited knowledge about the regulatory functions of the RNA molecules in teleost fish, including European seabass (Sarropoulou et al., 2025). Sarropoulou et al., 2025 demonstrated potential involvement of blood circulating miRNAs in modulating immune response against *Aeromonoas veronii* bv. *Sobria* in farmed European seabass.

Therefore our current study aims to characterize the association between brain miRNA expression and VNN susceptibility in farmed European seabass. Consequently, our study will; 1) expand the knowledge of the Europeans seabass miRNome; 2) identify miRNAs targeting the 3’ UTR of *ifi27l2a*, focusing on QTL-linked SNPs; 3) reveal associations between these SNPs in the 3’ UTR of *if27l2a* and miRNA binding/modulation to explain differential VNN resistance phenotypes (susceptible, intermediate, resistant).

## 2. Material and Methods

### 2.1 Animal production

Experimental fish were generated in January 2020 from a commercial broodstock confirmed free of NNV by PCR testing, from Valle Cà Zuliani Società srl (Pila di Porto Tolle, Rovigo, Italy). The details of the experimental population generation and challenge experiment have been described in our recent study that used the same animals (Mukiibi et al. 2025). Briefly, female fish underwent ovarian biopsies to select those at an appropriate egg development stage and those suitable dams received hormonal treatment with LH-Rha (luteinizing hormone releasing-hormone analogue, 10 µg/kg body weight) and were stripped 72 hours later. Eggs were promptly fertilized by mixing with milt from pre-collected sperm, then transferred to hatchery tanks (2000-L tanks with conical bases) after 5-6 minutes. A full-factorial mating design was implemented utilizing 25 dams and 25 sires, with fertilized eggs pooled and allocated across four rearing tanks. At 240 days post-hatch, and average weight of 13 g,, fish were transported to Istituto Zooprofilattico Speriimentale delle Venezie (IZSVe, Legnaro, Padova, Italy) where the disease challenge study was performed. A total 214 fish underwent VNN exposure to evaluate transcriptomic responses to infection. These fish received VNN challenge via intraperitoneal injection of 0.1 mL of a 1:100 diluted viral suspension (RGNNV 283.2009, stock titre 10^8.30^ TCID50 per mL). For each fish, brain tissue was collected at 48 h post-challenge for RNA sequencing, corresponding to the immune response peak.

### 2.2 Total RNA extraction

Basing on the major VNN QTL identified by Mukiibi et al. 2025, the 214 fish were grouped into resistant, intermediate and susceptible depending on their genotype for the QTL where by the resistant were homozygous to the resistant allele (RR), intermediate were heterozygote (RS) and susceptible were homozygote for susceptible allele (SS). Relatedly within each QTL genotype group the fish showed binary genetic variability mainly due to the genetic origin of their dams, hence the sample were grouped into two genetic clusters (refered to as cluster A and B in this study). Of the 214 fish, 21 fish were considered for miRNA profiling which included 9 samples (3 SS, 3 SR and 3 RR samples) from cluster A and 12 samples from cluster B (4 SS, 4 SR and 4 RR samples). Total RNA from the brain tissue was extracted using the Qiagen RNeasy Plus Universal Mini Kit (Qiagen, Toronto, ON, USA) according to the manufacturer’s instructions. A NanoDrop 2000 Spectrophotometer (Thermo Sciencitific, Wilmington, DE, USA) was used to quantify the RNA. We obtained total RNA with with absorbance ratios (A260/280) ranging between 1.8 and 2.0. RNA integrity number (RIN) values for all samples were higher than 8 which deemed them to be high quality and suitable for cDNA library preparation and downstream transcriptomic profiling.

### 2.3 Small RNA library construction and high throughput sequencing

A small RNA library was generated from the first sampling previously detailed, pooling equal concentrations of RNA from the different sample of brain. A small RNA library was generated from this pool a broad range of tissues and ages using the Illumina Truseq Small RNA Sample Preparation Kit (Illumina, San Diego, CA, USA) according to the manufacturer’s protocol. The purified cDNA library was sequenced in one Illumina Gallx platform run generating 40 bp single-end reads. Raw sequences were filtered eliminating adapter sequences, putative contamination and sequences between 15 and 32 bp were retained.

### 2.4 Bioinformatic sequencing data processing and miRNA expression profiling

Raw sequence reads were initially assessed for sequencing quality using FASTQC version 0.12.1 (Andrews, 2010). The reads were evaluated for quality based on multiple parameters such as average read length, adaptor content, per sequence GC content, and per base sequence quality scores. Thereafter, the Illumina universal adapter sequence (AGATCGGAAGAGCACACGTCTGAACTCCAGTCA) was trimmed from all the raw read sequences using Cutadapt V. 3.5 (Martin, 2011). In this study, reads of lengths shorter than 15 bp, or longer than 28 bp were removed as short or long reads, respectively. The retained reads were filtered for short RNA including ribosomal RNAs (rRNAs), transfer RNAs (tRNAs), small nuclear RNAs (snRNAS), and small nucleolar RNAs (snoRNAs), download from the Rfam database via RNA central a non-coding database (https://ftp.ebi.ac.uk/pub/databases/RNAcentral/releases/24.0/rfam/, accessed March, 2025). The final processed sequence reads were revaluated for quality using FASTQC.

To profile both novel and known miRNA expression in the samples from the cleaned sequenced data, the miRDeep2 package version 0.13 (Friedländer et al., 2012) was used together with the *Dicentrarchus labrax* genome assembly (version: GCA_905237075.1: dlabrax2021). The known seabass mature miRNA sequences and their precursor sequences from the miRbase database (release 22.1) and FishmiRNA database (Version September 2023) (Griffiths-Jones et al., 2008) (Desvignes et al., 2022), those of other teleosts: *Astatotilapia burtoni, Cheanocephalus aceratus, Oryzias latipes, Tetraodon nigroviridis, Fugu rubripes, Danio rerio, Cyprinus carpio, Gadus morhua, Hippoglossus hippoglossus, Paralichthys olivaceus, Ictalurus punctatus, Salmo salar, Petromyzon marinus, Gasterosteus aculeatus, Chaenocephalus aceratus, Perca fluviatilis*, as well as miRNAs from *Gorilla, Homo sapiens, Mus musculus*. The mapper module (mapper.pl) with default parameters was used to collapse reads of the sequences into unique read clusters, and then bowtie 1.3.0 short sequence aligner (Langmead et al., 2009) was employed to align the collapse reads to the indexed Dlabrax2021 reference genome. Using default parameters and input files including the reference genome, collapsed reads versus reference genome alignment, all known mature miRNAs and their precursor sequences (including the hairping structures). All known miRNAs were quantified by the miRDeep2 module (miRDeep2.pl), hence producing read counts for each individual sample. Subsequently, miRDeep2 was used to predict possible novel miRNAs and their respective precursors based from the read alignments to the seabass reference genome. Genomic regions stacked with aligned reads were excised as potential precursors and evaluated by the RNAfold tool within ViennaRNA-2.5.0 (Lorenz et al., 2011) for their potential to form stable secondary structures (hairpins), their ability to be partitioned into mature, loop and star strand, and their base pairing in the mature miRNA region. Subsequently, for each predicted miRNA, a mature miRNA consensus sequence, a precursor sequence, RNAfold *p-value*, the miRDeep2 score, and the probability that the predicted miRNA was a true positive were estimated and produced as outputs. Additionally, for each predicted novel miRNA, miRDeep2 produced their hairpin structure of the precursor sequence and the aligned read counts for each sample. The hairpin secondary structure and read counts by sample were also generated. Novel miRNAs were assigned provisional precursor IDs. Multiple mature sequences mapped to the same precursor, these unique mature sequences were distinguished by appending numerical suffixes to the provisional precursor IDs to avoid ambiguities in downstream analyses.

### 2.5 Differential miRNA expression analysis

Initially, counts for each mature miRNA from more than one precursor were averaged. Thereafter, all miRNAs that had less than 10 total read counts across the studied samples within each group were filtered out. Then miRNA expression variation patterns between samples in each group were visualized through a principle component analysis of the read counts using the DESeq2 Bioconductor package (Love et al., 2014) and the ggplot2 R package (Wickham, 2016). The differential expression analyses for both known and novel miRNAs was performed using the egdeR Bioconductor package in R (Robinson et al., 2009). To reduce false positive rates of the analyses, miRNAs within samples from each of the samples that had less than one count per million (CPM) in at least half of the analysed samples, were filtered out from the analyses, as proposed by Anders et al. 2013. For the retained miRNAs, their counts were normalized using the TMM method (Robinson & Oshlack, 2010). To test for differential miRNA expression between (SS, SR and SS) using the egdeR package in R. Normalized count data were modelled with a generalized linear model (GLM) under a negative binomial distribution to assess differences in each genetic cluster group between the three experimental groups: Resistant, Intermediate and Susceptible. Pairwise comparisons were conducted using specific contrasts within the GLM framework: Intermediate vs Resistant (with Resistant as the reference group), Susceptible vs Resistant (with Resistant as the reference group) and Susceptible vs Resistant (with Resistant as the reference group). For each comparison, miRNAs were considered differentially expressed if they exhibited a FDR <0.10 and an absolute log fold change greater than 1.5. Expression of the DEmiRNAs across samples between comparisons was visualized via hierarchical clustering heatmap plots using Pheatmap package (V.1.0.13) in R (Kolde, 2015).

### 2.6 miRNA Target prediction

Target gene prediction for the differentially expressed miRNAs (DEmiRNAs) from each pairwise comparison was performed using miRNAconsTarget from the sRNAtoolbox web server (https://arn.ugr.es/srnatoolbox/) (Rueda et al., 2015). Predictions were done on the 3’ UTR sequences of three candidate genes ENSDLAG00005026832, LOC127358630 and LOC127358685 previously identified as the main modulators of VNN resistance in European seabass (Mukiibi et al., 2025). These regions were used for DEmiRNAs target prediction with animal-based tools: TargetSpy 1.0 (Sturm et al., 2010), miRanda v3.3a (Enright et al., 2003) and PITA (Kertesz et al., 2007) with default parameters. To increase reliability, TargetScanFish 6.2 (https://www.targetscan.org/fish_62/) (Lewis et al., 2005) as was described by Dewaele et al. 2025, and IntaRNA V. 2.0 (Mann et al., 2017), were also used with default parameters and the same 3’UTR-database. For filtering of target predictions, binding affinity thresholds were set to minimum free energy (MFE) ≤ -25.0 and/or score ≥ 150 for miRanda; MFE ≤ - 18 for TargetSpy with score of ≥ 0.998; MFE ≤ -15.0 for PITA; seed match types restricted to 8mer-1a or 7mer-m8 for TargetScan and Context+ Score ≤ -15.0; and IntaRNA with energy ≤ -9 and Hybridization Energy ≤ -20.0. 3’ UTRs with perfect complementary base-pairing to the miRNA seed region (base pairs 2-7) were taken into consideration. Only gene targets predicted by at least three tools were retained for downstream analysis.

## 3. Results

### 3.1 MiRNA sequence data and alignment quality

On average the Illumina next generation sequencing yielded over 18.0 M (million), 15.6 M and 17.8 M high quality raw reads per sample for susceptible, intermediate and resistance to VNN resistance QTL genotypes samples (Table S1). After 3’ adaptor clipping, an average of 22.16%, 15.41% and 30.64% of the reads were removed as long reads (>28bp), and an average of 9.72%, 14.04% and 8.89% of the reads were removed as short reads (<15bp) of SS, SR and RR samples (Table S1). Additionally, on average 50.94% of the reads were removed since they aligned to 35.29% rRNAs, 7.9% tRNAs, 4.14 % snRNAs and 1.6% snoRNAs, respectively (Table S1). An average of 6.0 M reads were retained for miRNA profiling analysis by mirDeep2 (Table S1). Most of the reads ranged between 20 and 24 bp in length as shown in Fig 1A, with an average length of 22 bp. The retained reads were of high quality as depicted by high average Phred scores in Fig. 1B. Of the retained reads mapped to the Dlabrax2021 reference genome on average, ranging from 35.15%, 39.91% and 25.79% for SS, SR and RR (Table S1).

**Fig 1.**
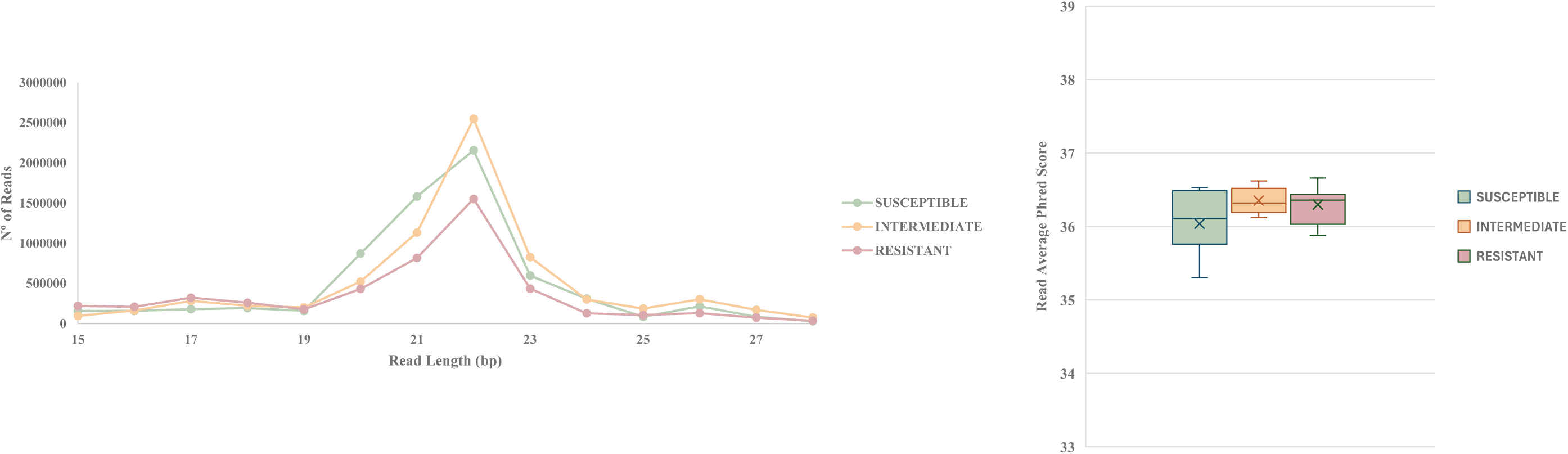
(**a**) Line plot showing the read length distributions in the final cleaned sequence data after quality control involving 3′ Illumina sequencing adaptor clipping, removing very long reads (> 28 bp) and short reads (< 15 bp), and removing reads that mapped to other small RNA species (rRNAs, snRNAs, tRNAs and snoRNAs) for Susceptible, Intermediate and Resistant samples for all samples. (**b**) Box plots showing the average Phred quality scores of the retained reads for all samples.

### 3.2 Genetic clusters

Population structure analysis from Mukiibi et al. (2025) identified two distinct genetic clusters (A and B) among VNN-challenged dam fish, likely representing different geographic origins with opposite phenotypic variation in VNN resistance. From this dataset, 21 dams were specifically selected for the present study: cluster A containing 9 and cluster B comprising 12 individuals (Table S2).

### 3.3 Known miRNA expression and novel miRNA profiles

Cluster A revealed 304, 284 and 212 expressed known miRNA in SS, SR and RR samples, respectively. Of all these, 139 unique known miRNAs (34%) were common to all the three susceptibility groups to VNN as shown in Fig 2A. Regarding to cluster B we identified 331, 326 and 351 expressed known miRNA in SS, SR and RR samples, respectively. Of all these, 212 unique known miRNAs (45%) were common to all the three susceptibility groups as shown in Fig 2C. All expressed known miRNAs identified, and their average aligned read counts in each group are presented in Table S3.

**Fig. 2.**
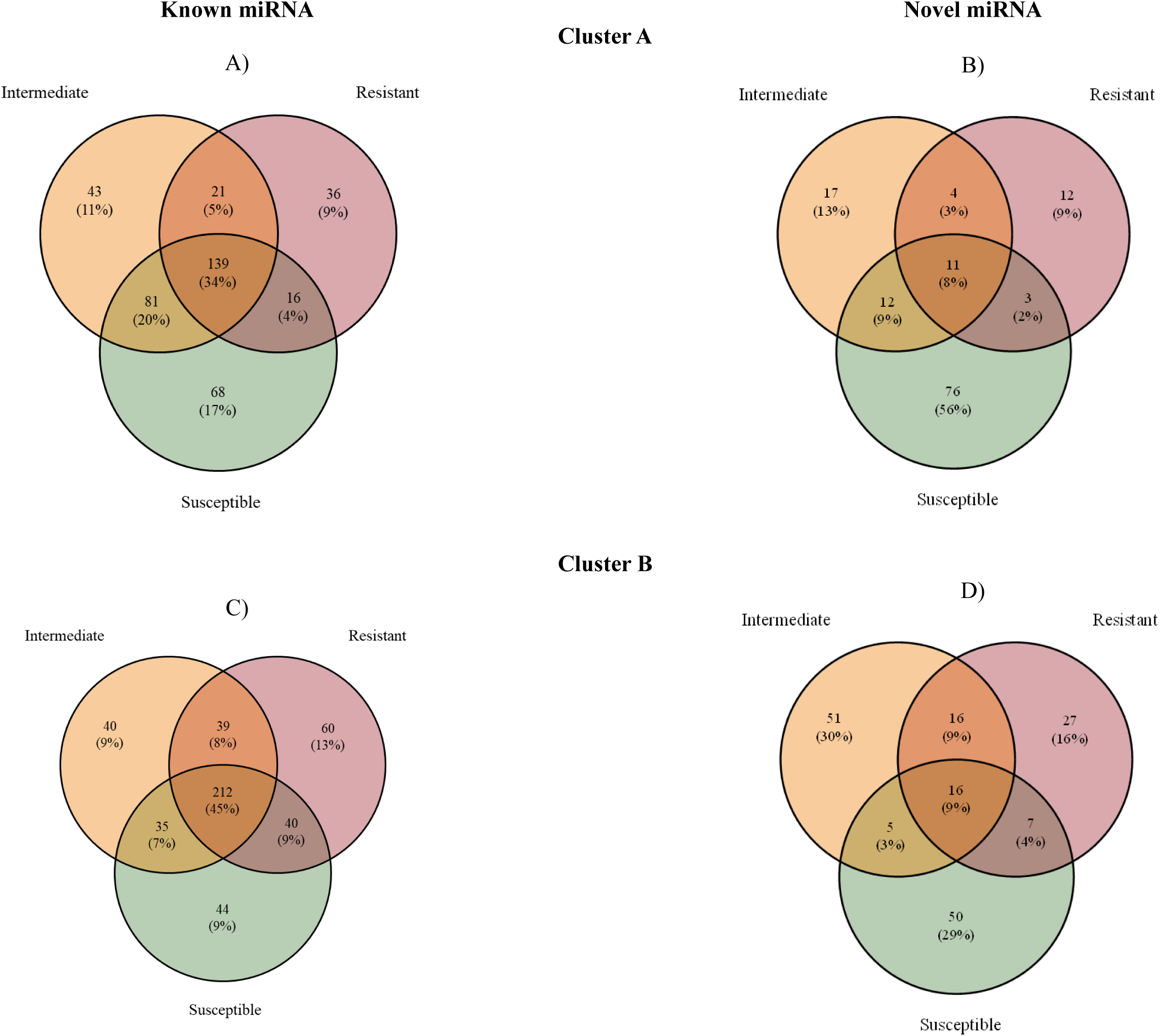
Venn diagram showing overlap of expressed known (A, C) and novel (B, D) miRNAs in the brain tissue of seabass from the three studied groups (Intermediate, Susceptible, and Resistant) in cluster A and B, respectively.

Novel miRNA prediction (mirDeep2 score ≥4, an estimated probability that the predicted miRNA candidate is a true positive is greater than 70%, and with a significant Randfold p-value suggesting that the miRNA’s precursor sequence could be folded into a thermodynamically stable hairpin), identified 102 (from 124 precursors), 44 (from 51 precursors) and 30 (from 43 precursors) novel miRNAs in SS, SR and RR samples respectively in cluster A (Fig 2B). Meanwhile, in cluster B, we identified 78 (from 97 precursors), 88 (from 116 precursors) and 66 (from 81 precursors) novel miRNAs in SS, SR and RR samples respectively (Fig. 2D). All the identified novel miRNAs and their miRDepp2 prediction scores are provided in Table S4.

### 3.4 miRNA differential expression

Differential miRNA expression analysis was performed in cluster A and B separately. To each cluster different comparisons were performed between susceptible *vs* resistant, susceptible *vs* intermediate and intermediate *vs* resistant. At a fold change > 1.5, and FDR < 0.10, in cluster A the comparison between SS*vs*RR identified 34 DE known miRNAs in the brain tissue, in which 30 were upregulated and 4 downregulated in susceptible group against resistant (Table S5A, Fig 3A). However, SS*vs*SR and SR*vs*RR did not shown significant DEmiRNAs (Fig. 3A). In cluster B, 4 DE miRNAs including 3 known miRNAs and one novel miRNAs were identified in SS*vs*RR, with all DE miRNAs upregulated in susceptible group regarding resistant seabass(Table S5B, Fig 3B). In the comparison SS*vs*SR three DE miRNAs, with two known miRNAs and one novel, all showing upregulation in susceptible regarding intermediate samples (Table S5C, Fig 3B). And in the last comparison of intermediate *versus* resistant group, it was found two known miRNAs DE upregulated in intermediate samples respect resistant samples, meanwhile two DE miRNAs (one known and another novel) were downregulated in the samples group regarding resistant (Table S5D, Fig. 3B). Considering all the miRNA differential expression analyses performed, DEmiRNAs identified varied greatly among the comparisons, as shown in the UpSet plot (Fig. 4). Notably, miR-199-5p was the common miRNA shared between Cluster A and B interactions, upregulated in susceptible samples. Within Cluster B, the novel DEmiRNA, NW_026136709.1.1 was common between SS *vs* SR and SS *vs* RR (both upregulated in susceptible samples). Finally, miR-96-5p (SS *vs* SR and SR *vs* RR), exhibited downregulation in intermediate samples (Table S5).

**Fig. 3.**
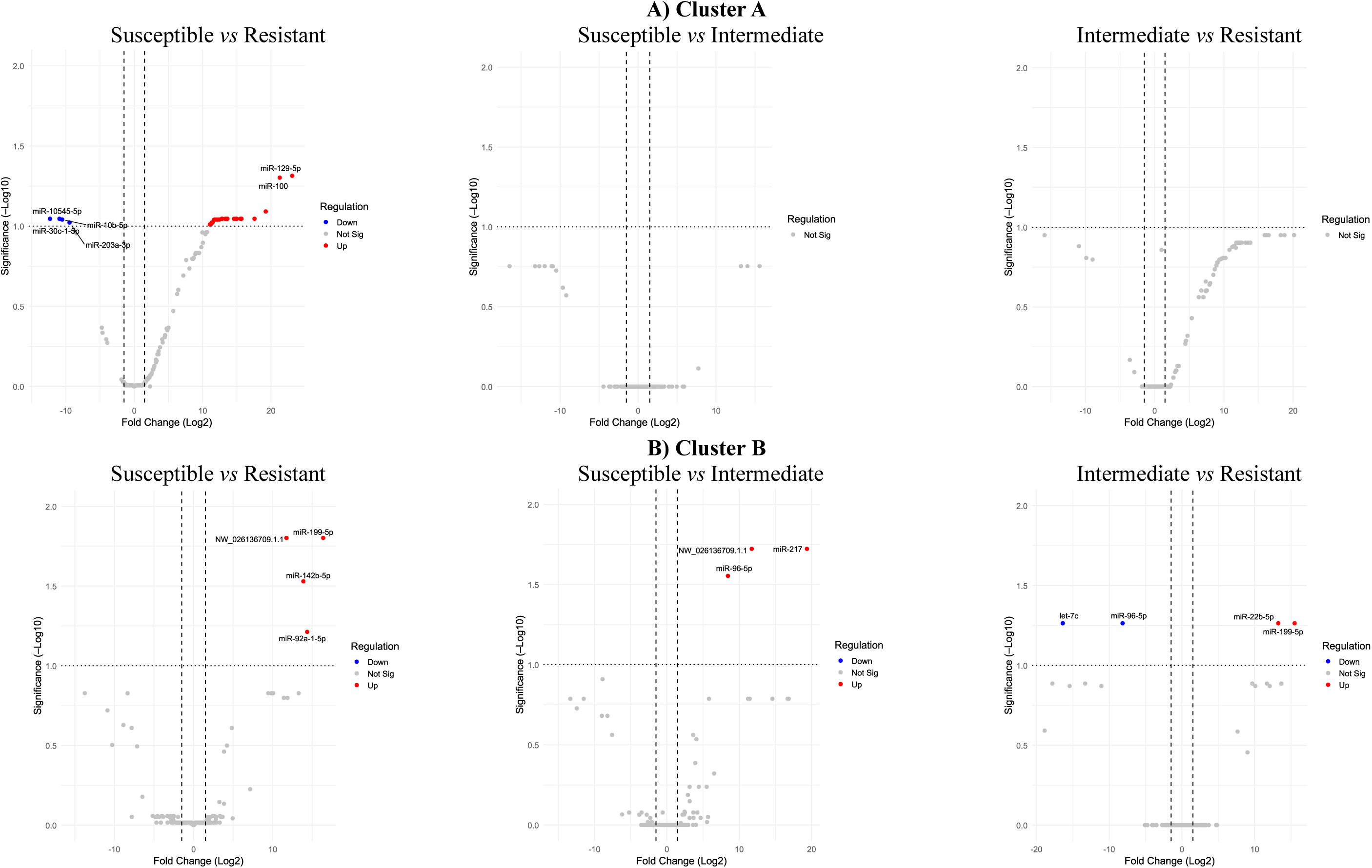
Volcano plot of differentially expressed miRNAs in the brain tissue of seabass from the three studied [Intermediate, Susceptible, and Resistant] in cluster A (A) and in cluster B (B).

**Fig 4.**
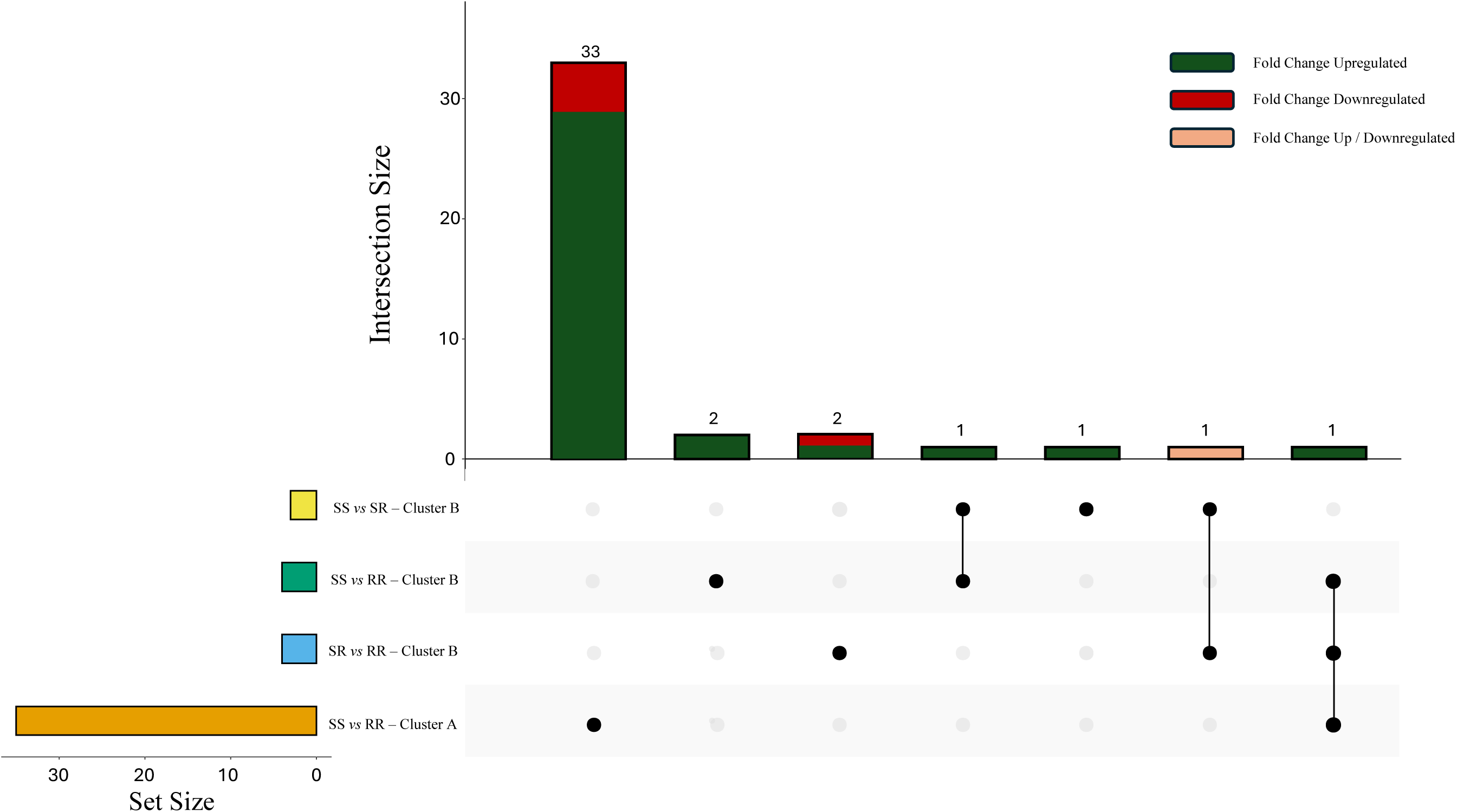
UpSet plot illustrating the overlap of DEmiRNAs across comparisons. Vertical bar height represents the number of DEmiRNAs shared among specific combinations of samples. Horizontal bar widths represent the number of DEmiRNAs specific of each comparison. Yellow indicates susceptible (SS) *vs* Intermediate (SR) in cluster B, green SS *vs* Resistant (RR) in cluster B, clear blue SR *vs* RR, and orange SS *vs* SR in cluster A. Coloured set size bars and interactions dots correspond to each comparison. Numbers above each bar indicate the intersection size of DEmiRNAs. Bar colours indicate regulation status, green shows upregulated DEmiRNAs (Fold Change ≥ 1.5 in one or more comparisons), red shows downregulated DEmiRNAs (FC ≤ 1.5 in one or more comparisons) and orange indicated discordant regulation between comparisons in each DEmiRNAs (up/down across comparisons).

In addition, heatmaps were generated by clustering miRNA expression across all samples using Euclidean distance. Both susceptible and resistant samples showed cluster-specific patterns consistent across genetic clusters A and B. Notably, heatmaps revealed consistently higher miRNA expression in susceptible *versus* resistant samples (Fig 5). This trend is conserved in cluster B for the DEmiRNA for the Susceptible *vs* intermediate comparison, with more expression in susceptible samples (Fig. S1A). In contrast, the intermediate *versus* resistant group exhibited clear group separation despite some sample overlap (Fig. S1B).

**Fig 5.**
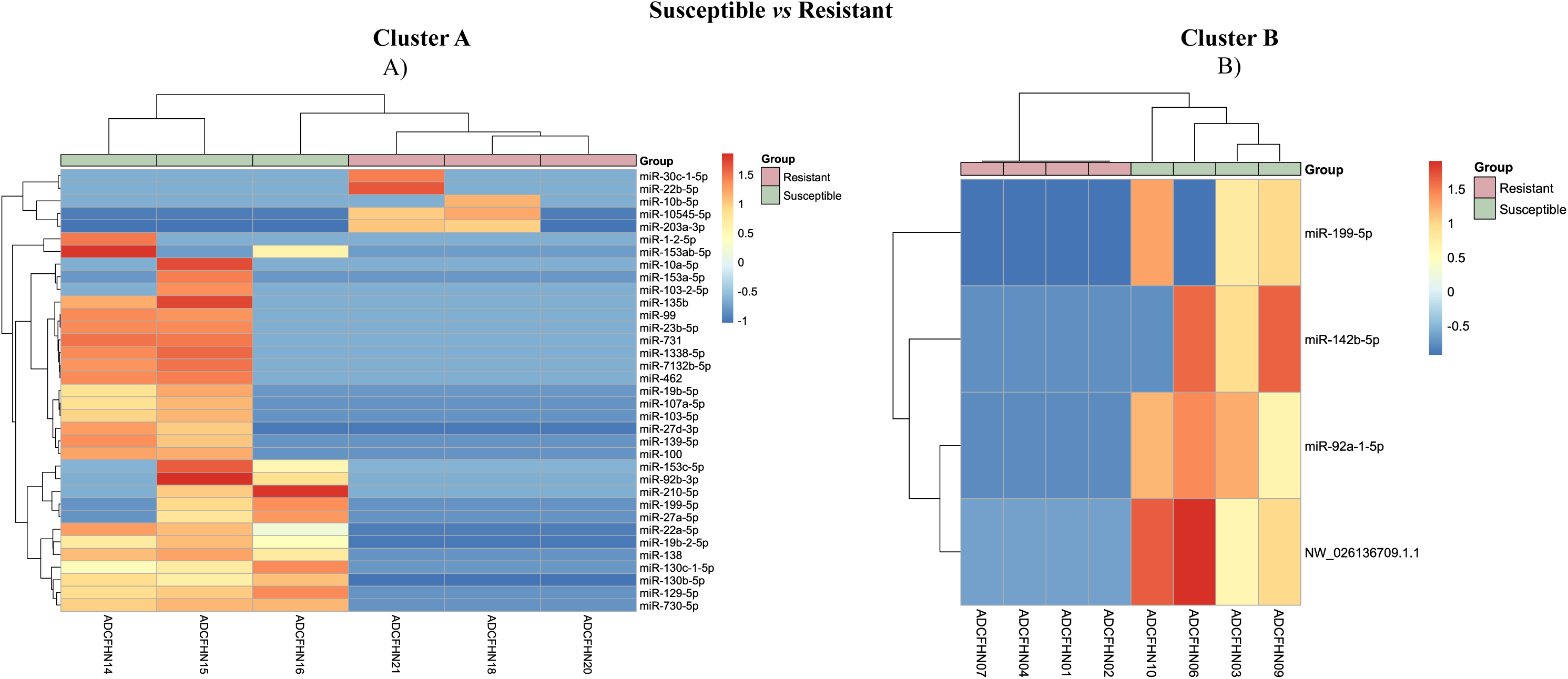
Heatmap of differentially expressed miRNAs between susceptible and resistant samples in cluster A (A) and in cluster B(B). Colour scale: blue (low expression) to red (high expression). Sample groups: green (susceptible) and pale pink (resistant).

### 3.5 Target gene prediction for DEmiRNAs

Target gene prediction was performed for the differentially expressed known and novel miRNAs identified across comparison (Table S5). This analysis focused on finding potential regions target for the interferon-induced genes (LOC127358685 (*ifi27l2*), LOC127358630 (*ifi27l2a*), and ENSDLAG00005026832 (*ifi27l2a*)), which have been reported as containing SNPs for the major QTL for resistance to VNN (Mukiibi et al., 2025). We employed a multi-tool consensus approach, as described in the “Material and Methods” section, retaining only those targets predicted by at least in three out of the five tools. Target prediction identified one miRNA with two different significant targets using miRanda, 8 miRNAs with PITA, 12 miRNAs with TargetSpy, 4 miRNAs with TargetScan, and 2 miRNAs with IntaRNA (Table S6). Among the differentially expressed miRNAs, miR-199-5p uniquely exhibited statistically significant interactions with ENSDLAG0005026823 across all five prediction tools, targeting two distinct miRNA Recognition Elements (MREs) (Fig. 6; Table S6). The first MRE (Chr3:10,082,340-10,082,359) was predicted with high confidence by miRanda (score = 154, MFE = - 14.29), TargetSpy (score = 0.999, MFE -13.30), and TargetScan (7mer-m8, context++ score = - 0.178). The second MRE (Chr3:10,082,403-10,082,425) showed the strongest biding energy, as predicted by Miranda (MFE = -25.06, score = 142), PITA (MFE= -19.4), and IntaRNA (Total energy = -9.37 and Hybridization Energy = -20.64) (Table S6; Fig 6). Notably, both MREs flank a VNN resistance-associated SNP (Chr3:10,082,380) (Mukiibi et al., 2025), with a distances of 22 bp and 39 bp, respectively, between the SNP and the seed regions of MRE1 and MRE2.

**Fig. 6.**
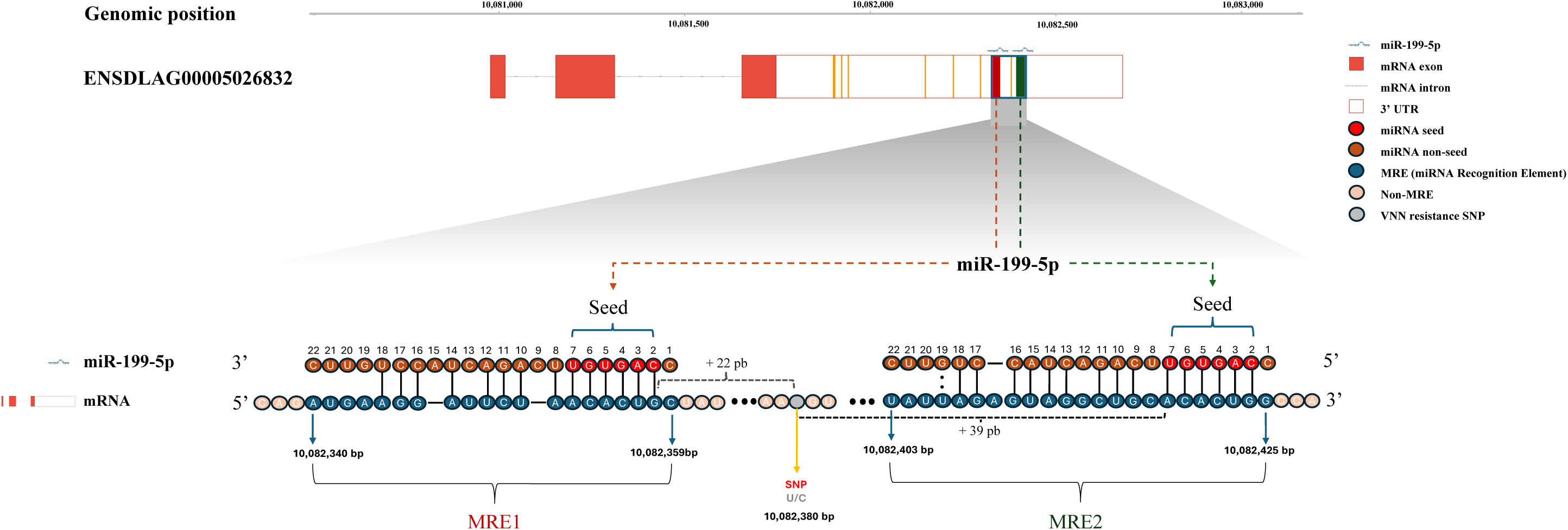
Genomic landscape of miR-199-5p target sites in ENSDLAG00005026832 (*ifi27l2a*) on chromosome 3. The figure shows the ENSDLAG00005026832 gene with miRNA target sites (blue boxes) containing two miRNA Recognition Elements (MREs): MRE1 (dark red) and MRE2 (dark green), adapted from miRanda-predictions. The zoomed region illustrates the miRNA seed region (red nucleotides), miRNA non-seed region (brown nucleotides), Watson-Crick base pairing (solid lines, |), G:U Wobble pairs (dotted lines, :), MRE nucleotides (dark blue), non-MRE (pale pink), and VNN resistance-associated SNP (grey).

### 3.6 Correlation of miR-199-5p and mRNAs expression in brain tissue

To determine if miR-199-5p have a direct influence on resistance to VNN, we conducted the Spearman’s correlation for interactions between the expression of miR-199-5p and mRNA expression for cluster A and Cluster B (data RNAseq data for sample samples obtained from Mukiibi et al., 2025). Significant negative correlation was found only in cluster A (*p*-value= 0.002, r = - 0.840) (Fig. 7A), but not in cluster B (*p*-value = 0.927, r = 0.448) (Fig. S2a). Notably, we observed lower miR-199-5p expression in VNN resistant samples for cluster A (Fig. 7B) and a similar pattern was shown in the boxplot of Cluster B (Fig. S2B), contrasting with the higher expression of *ifi27l2/2a* mRNA in resistant seabass (Mukiibi et al., 2025).

**Fig 7.**
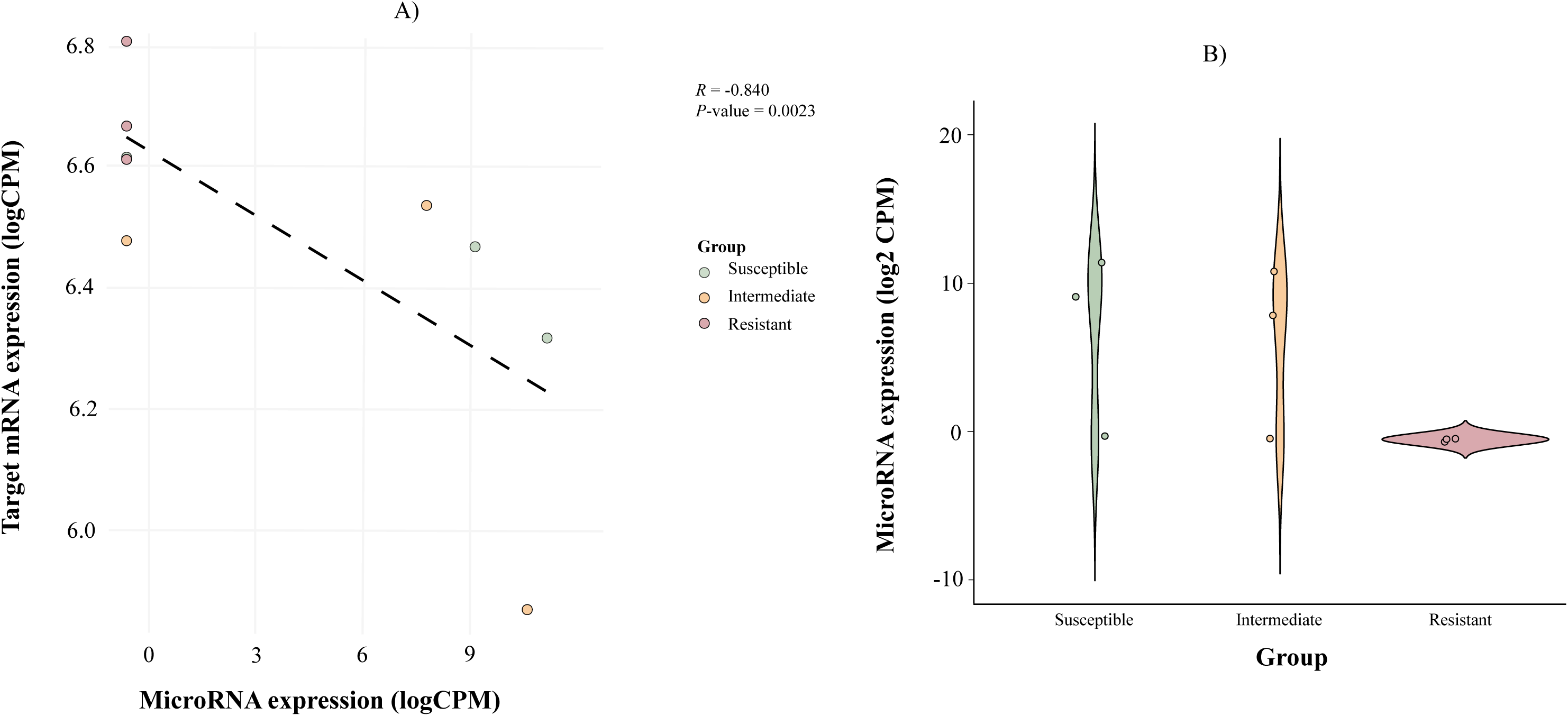
A) Spearman’s correlation between the expression of miR-199-5p and the target gene ENSDLAG00005026832 (*ifi27l2a*) (P = 0.0023, r =-0.840) in Cluster A. B) Violin plot showing the gene expression of miR-199-5p in animals with different VNN resistance QTL genotypes

## 4. DISCUSSION

MicroRNAs (miRNAs) are short (∼20–22 nucleotides), highly conserved post-transcriptional regulators that typically repress targets gene by binding complementary sequences in mRNA 3’ UTRs (triggering mRNA degradation or translational repression), though less commonly stabilizing mRNAs or targeting 5’ UTRs or promoters (Bartel, 2009). In teleosts, miRNAs orchestrate diverse developmental processes, environmental adaptation, and robust immune and inflammatory responses to pathogens, including bacteria (e.g., *Aeromonas, Vibrio*) and viruses such as siniperca chuatsi rhabdovirus or orthomyxovirus (Andreassen & Høyheim, 2017; Bizuayehu & Babiak, 2014; Zhou et al., 2023). However, the brain miRNome during VNN infection, and its crosstalk with the major chromosome 3 resistance QTL remains uncharacterised in European seabass, a critical gap, given that the brain is a primary target organ for VNN. In this study, we identified brain miRNAs across susceptible, intermediate, and resistant fish defined by the chromosome 3 QTL (Mukiibi et al., 2025), revealing the post-transcriptional networks underlying VNN resistance.

Small–RNA sequencing generated high–quality libraries, retaining approximately 6 million post–filtered reads per sample with a clear 22-nucleotide peak and high Phred scores, consistent with established standards for robust miRNA discovery and quantification (O’Brien et al., 2018; Papadaki et al., 2022; Sarropoulou et al., 2025; Sun et al., 2014). In addition, population structure analysis identified two genetic clusters likely from distinct geographic origin among dams, which were analyzed separately to control background effects geographic origins (Barsøe et al., 2021; Mukiibi et al., 2025; Vela-Avitúa et al., 2022). This approach revealed two key patterns, conserved and cluster-specific regulatory patterns, revealing a shared core of known miRNAs across genotypes supporting baseline homeostasis alongside 56% more novel miRNAs in cluster A susceptible fish. This was in concordance with the observed in IPNV-susceptible Atlantic salmon (Woldemariam et al., 2020).

Differential expression analyses revealed marked miRNA dysregulation in susceptible fish, particularly in Cluster A, where SS genotypes showed substantially higher levels of upregulated DEmiRNAs than RR fish. This pattern is in accordance with a study reported in Atlantic salmon challenged with IPNV, where susceptible samples with high viral loads showed stronger miRNA upregulation than resistant, which controlled viral replication (Woldemariam et al., 2020). Therefore, these results suggest an infection-induced, hyperactive miRNA response in genetically seabass that is ultimately maladaptive, potentially suppressing key antiviral effectors in line with pro-viral miRNA activities reported in other teleost species (Samir et al., 2016; Zhang et al., 2017, 2021) .

### 4.1 MiR-199-5p: Central regulator of ifi27l2a

Our results identify miR-199-5p as a pivotal post-transcriptional regulator *ifil27l2a*, gene associated with the major VNN resistance QTL on chromosome 3 in seabass. Recent functional-genomic work found that this QTL interval contains multiple copies of *ifi27l2a* and one copy of *ifi27l2*, thereby explaining a significant component of the 90% survival advantage conferred by favourable QTL alleles (Mukiibi et al., 2025). In particular, *ifi27l2a* encodes for interferon alpha-inducible inducible protein, which is an interferon-stimulated gene (ISG) that are activated following IFN signalling and contribute to antiviral defence by limiting replication and promoting viral degradation (Schoggins, 2018). In mammals, *ifi27l2a* exhibits antiviral properties in the central nervous system, restricting virus replication (Lucas et al., 2016; Schoggins, 2018). In zebrafish, *ifi27l2a* homologues supresses viral replication by targeting viral phosphoprotein, suggesting a conserved antiviral role in teleost (Guo et al., 2023). In European seabass and other teleosts, this gene is induced by VNN (Nuñez-Ortiz et al., 2016; Toubanaki et al., 2022). By mapping miRNA-mRNA interactions onto this locus, our study adds a post-transcriptional layer to the model in which miR-199-5p could be a direct negative regulator of *ifi27l2a* in the brain tissue.

The pivotal role of miR-199-5p is supported by the following converging lines of evidence I) It is the only DEmiRNA shared between both genetic clusters (A and B) in SS *vs* RR, with consistent overexpression in susceptible fish. II) It remains significant under stringent FDR threshold in cluster B. III) MiR–199–5p is the only microRNA that shows consensus binding (≥3/5 tools) to two MREs within the 3’ UTR of *ifi27l2a*. IV) It exhibits a strong inverse correlation with *ifi27l2a* mRNA. Taken together, these results strongly suggest allele and background dependent repression of *ifi27l2a* by miR-199-5p.

At the expression level, miR–199–5p was consistently upregulated in SS *versus* RR fish across both genetic clusters, directly supporting the first two lines of evidence. Its persistence as a significant DEmiRNA under stringent FDR control in the cluster B background further reinforces the robustness of this signal. The fact that this miRNA is the only DEmiRNA targeting the immune gene *ifi27l2a* is supported by extensive cross-species evidence revealing pleiotropic immune roles. In human oral submucous fibrosis, miR-199-5p overexpression in buccal fibroblasts induces apoptosis (Yuan et al., 2019). In dairy cattle, milk exosomal miR-199-5p is enriched during subclinical mastitis, with predicted targets in immune and inflammatory pathways (Mahala et al., 2024). In yellow catfish, a teleost model, miR-199-5p directly represses mTOR, suppressing the autophagy-mediated stress response during ammonia toxicity (Wang et al., 2024). Collectively, across mammals and teleosts, miR-199-5p emerges as a conserved immune regulator of apoptosis, inflammation and stress adaptive pathways, aligning with a role as negative regulator of IFN-driven ISGs at the seabass VNN-resistance locus.

In addition, two high-confidence miR-199-5p recognition elements were identified in the *ifi27l2a* 3’ UTR: MRE1 (Chr3:10,082,340–10,082,359) and MRE2 (Chr3:10,082,403–10,082,425), flanking a candidate causal SNP (Chr3:10,082,380) at distances of 22 bp and 39 bp, respectively. This is notable because this region harbours the SNPs most strongly associated with VNN survival (Mukiibi, et al., 2025). Both MREs were supported by five independent prediction tools with favourable binding energies (Table S6, Fig. 7). Although the 22-39 bp spacing exceeds the canonical 13-35 nucleotides cooperativity window (Sætrom et al., 2007), long range cooperation can still occur through 3’ UTR secondary structure and protein scaffold interactions (Briskin et al., 2020).

Critically, SNPs flanking MREs can modulate binding affinity and repression efficacy, even when seed complementary remains intact, by altering local 3 UTR folding and RISC accessibility (Grimson et al., 2007; Rykova et al., 2022). In our research, resistant fish carry QTL alleles associated with higher survival and higher *ifi27l2a* expression, whereas susceptible genotypes show both increased miR-199-5p abundance and lower *ifi27l2a* levels. Rather than a complete disruption of miRNA-mRNA assembly, we therefore propose that allele-dependent differences in the 3’UTR context may adapt miR-199-5p binding efficiency and repression, acting together with differential miRNA expression. This allele-dependent modulation of miRNA targeting is consistent with broader frameworks in which 3’UTR SNPs function as cis-regulators of miRNA-mRNA interactions (Grimson et al., 2007; Rykova et al., 2022; Võsa et al., 2015).

Although miR-199-5p was consistently elevated in susceptible seabass across both clusters, its functional impact differed markedly. In cluster A, the strong negative correlation indicates effective miRNA-mRNA *ifi27l2a* repression, with susceptible fish lacking robust regulatory mechanisms. In cluster B, the absence of significant correlation, despite conserved miR-199-5p upregulation, suggests attenuation by favourable cis-regulatory variants (Mukiibi et al., 2025) and additional buffering layers, such as alternative polyadenylation generating resistant 3’ UTR isoforms (Mayr, 2016) or competitive endogenous (ceRNA) networks sequestering miR-199-5p (Salmena et al., 2011)), can attenuate miRNA dependence in the cluster B backgrounds (Zhang et al., 2012). Thus, miR-199-5p expression marks susceptibility across both populations, but functional target repression is most pronounced in the cluster A, revealing a multi–layered regulatory model of *ifi27l2a* locus.

Together, these observations support a model in which miR-199-5p acts as a negative regulator of the IFN-ISG axis at *ifi27l2a*. Susceptible seabass show higher miR-199-5p expression and lower *ifi27l2a* levels, consistent with enhanced post-transcriptional repression and reduced antiviral protein availability in neurons, thereby facilitating NNV replication. In contrast, resistant fish display lower miR-199-5p expression and higher *ifi27l2a* abundance, in line with a more permissive IFN-ISG response and increased survival. Because resistant and susceptible fish carry different alleles at the same chromosome 3 QTL, we hypothesize that alternative 3’UTR haplotypes at *ifi27l2a* may fine–tune miR–199–5p binding efficiency rather than completely disrupting miRNA-mRNA assembly. In susceptible haplotypes, the local 3’UTR context could favour more effective miR–199–5p recruitment and stronger repression, whereas resistant haplotypes may weaken this interaction, thereby contributing to higher *ifi27l2a* expression in combination with reduced miR–199–5p abundance. In resistant fish, this configuration could effectively releases *ifi27l2a* from post-transcriptional repression, enabling a more robust antiviral response and is consistent with the ∼90% survival advantage associated with this QTL (Mukiibi et al., 2025) This allele–dependent modulation of miRNA targeting is consistent with 3’UTR SNPs acting as cis–regulators of miRNA-mRNA interactions (Grimson et al., 2007; Rykova et al., 2022; Võsa et al., 2015).. In conclusion, our results identify miR–199–5p as a promising biomarker for VNN resistance, although functional assays will be required to validate the proposed regulatory mechanism.

## CONCLUSION

This work provides the first comprehensive characterization of the brain miRNome in European seabass under VNN challenge, revealing novel insights into the post–transcriptional mechanisms potentially acting at the major resistance QTL on chromosome 3. By integrating small RNA sequencing with genotype–specific comparisons across two genetically distinct populations, our findings demonstrate that both miRNA expression patterns and their regulatory impact are strongly shaped by population structure. This highlights the need to account for background genomic variation when interpreting immune regulation in aquaculture species. Susceptible fish exhibit consistent miRNA upregulation across genetic clusters, indicating broader immune suppression that genetic markers alone cannot capture. The identification of two MREs within the *ifi27l2a* 3’ UTR (located beside resistance–associated SNPs) provides new evidence that cis–regulatory variants may modulate miRNA accessibility and contribute to differences in antiviral gene expression. The genotype–dependent behaviour of these interactions suggests that variation at the QTL influences the degree to which miRNA repression can occur, supporting a regulatory model where VNN resistance emerges from the combined effect of favourable alleles and reduced miRNA-mediated inhibition. These insights advocate for miR-199-5p as a candidate biomarker for epigenetic-assisted breeding, bridging QTL selection with non-invasive monitoring to control aquaculture losses. Altogether, these findings reveal an additional regulatory layer at the VNN resistance locus and position miRNA–UTR interactions as relevant components of disease resistance in seabass. Future functional validation of these interactions will be essential to determine their causal role and to evaluate the potential integration of miRNA–associated markers into selective breeding strategies aimed at enhancing resistance in aquaculture populations.

## DATA ABAILABILITY

Genotypes and phenotypes have been submitted to the Europeans Variant Archive (EVA). Whole-genome raw sequencing data have been uploaded to the NCBI Short Read Archive (SRA) under BioProject accession PRJNA1110973 (https://www.ncbi.nlm.nih.gov/bioproject/PRJNA1110973/). RNA-sequencing data have also been uploaded to the NCBI SRA under BioProject accession PRJNA1122486 (https://www.ncbi.nlm.nih.gov/bioproject/PRJNA1122486/).

Additional data will be made available on request. Supplementary data has been attached.

## ACKNOWLEDGEMENTS

We thank Valle Cà Zuliani Società Agricola srl for providing the fish used for the VNN challenge experiment.

## FUNDING

This work was funded by the European Union’s Horizon 2020 research and innovation programme under grant agreement No 817923 (AQUAFAANG).). It also supported by postdoctoral programme of the Xunta de Galicia (Consellería de Cultura, Educación, Formación Profesional e Universidades) to R. R-V (ED481B-2023-104). DR was supported by BBSRC Institute Strategic Grants to the Roslin Institute (BBS/E/20002172, BBS/E/D/30002275, BBS/E/D/10002070 and BBS/E/RL/230002A), and by the Oportunius programme of the Axencia Galega the Innovación (GAIN, Xunta de Galicia).

### Credit Authorship contribution statement

R.R.V.: Conceptualization, methodology, software, formal analysis, investigation, data curation, visualization, writing - original draft, writing - review & editing; R.M.: Conceptualization, methodology, investigation, supervision, writing - review & editing ; S. F.: methodology, investigation, resources, writing - review & editing; R.F.: methodology, data curation, resources, writing - review & editing; L P.: methodology, investigation, resources, writing - review & editing; G. D. R.: methodology, investigation, resources, writing - review & editing; J.R.: methodology, data curation, resources, writing - review & editing; ; M.B.: methodology, investigation, resources, writing - review & editing; D.B.: methodology, data curation, resources, writing - review & editing; A.T.: methodology, data curation, resources; writing - review & editing; F. P.: methodology, data curation, resources; writing - review & editing; C. P.: methodology, investigation, resources, writing - review & editing; R.D.H.: funding acquisition, methodology, resources, project administration writing - review & editing; C.S.T.: funding acquisition, methodology, resources, project administration, writing - review & editing; L.B.: funding acquisition, methodology, resources, project administration, writing - review & editing; D.R.: methodology, investigation, supervision, resources, funding acquisition, project administration, writing - review & editing;

