## Supplementary material for "MicroRNA modulation of viral nervous necrosis resistance in European seabass": Fig. s1

### Cluster B

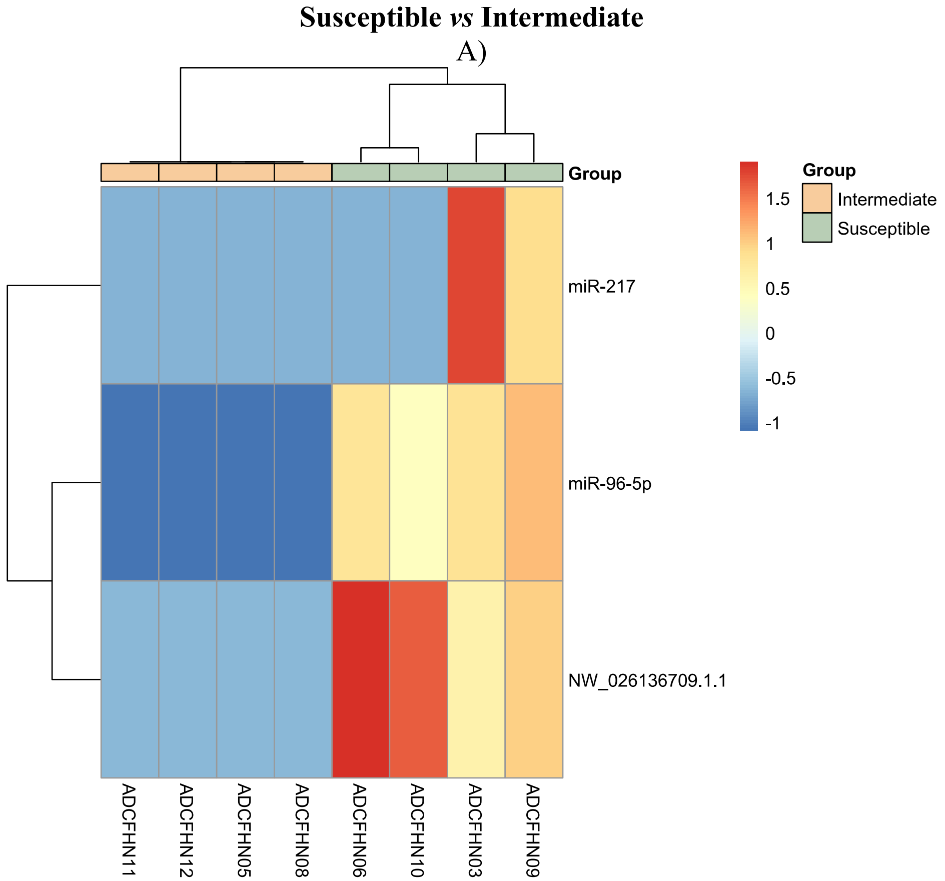

Susceptible *vs* Intermediate
A)
Group
miR-217
miR-96-5p
NW_026136709.1.1
ADCFHN11
ADCFHN12
ADCFHN05
ADCFHN08
ADCFHN06
ADCFHN10
ADCFHN03
ADCFHN09
1.5
1
0.5
0
-0.5
-1
Group
Intermediate
Susceptible


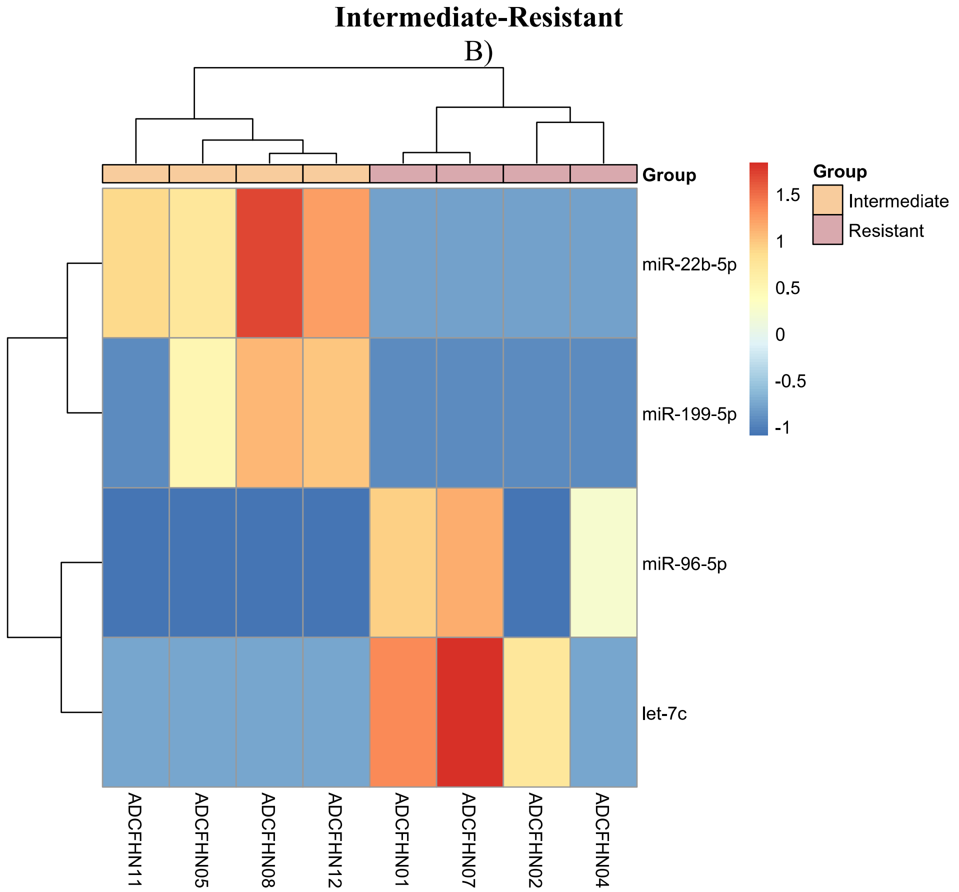

Intermediate-Resistant
B)
Group
miR-22b-5p
miR-199-5p
miR-96-5p
let-7c
ADCFHN11
ADCFHN05
ADCFHN08
ADCFHN12
ADCFHN01
ADCFHN07
ADCFHN02
ADCFHN04
1.5
1
0.5
0
-0.5
-1
Group
Intermediate
Resistant

**Fig S1.** Heatmap of differentially expressed miRNAs between susceptible and intermediate samples in cluster B (A) and intermediate and resistant samples in cluster B (B). Colour scale: blue (low expression) to red (high expression). Sample groups: green (susceptible), orange (intermediate) and pale pink (resistant).
