## Supplementary material for "MicroRNA modulation of viral nervous necrosis resistance in European seabass": Fig. S2

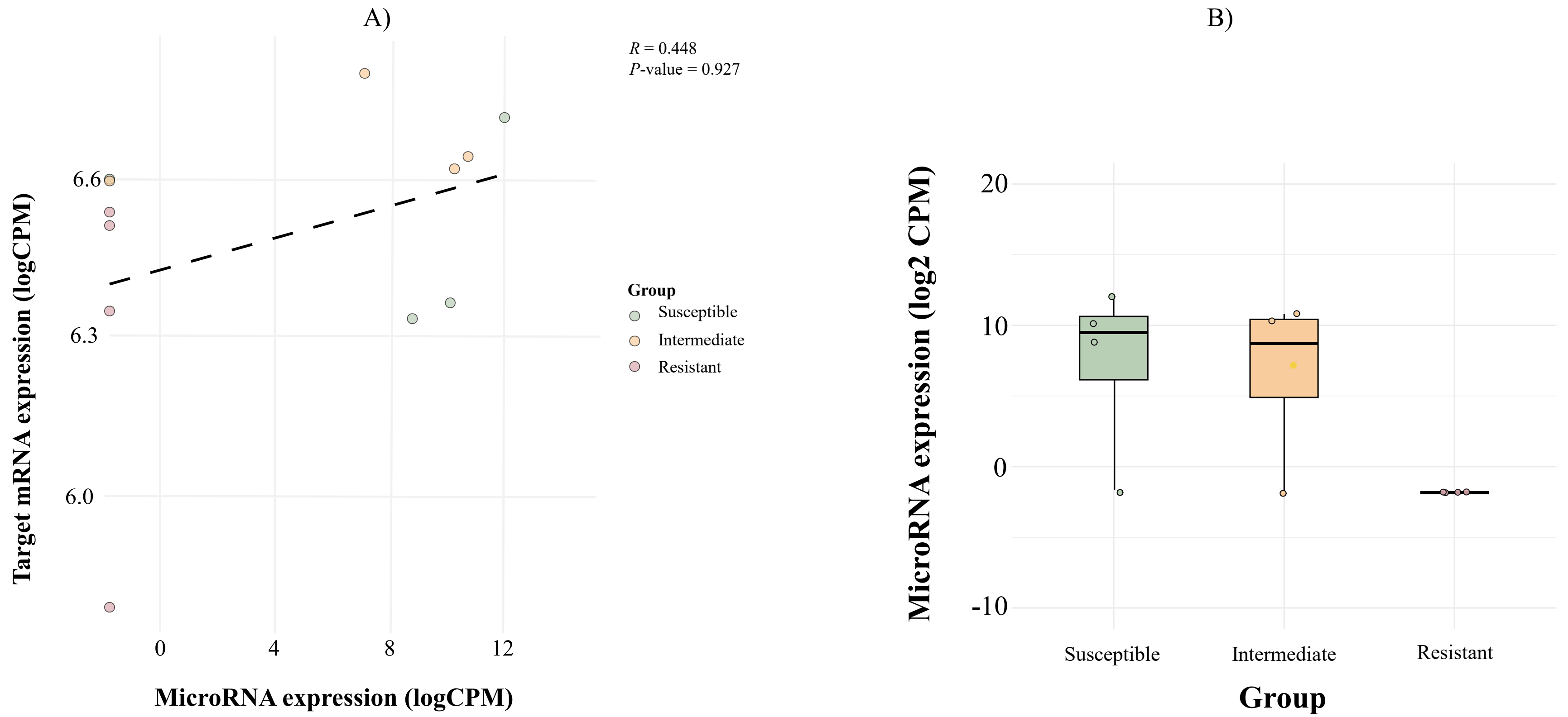


**Fig S2.** Spearman's correlation between the expression of miR-199-5p and the target gene ENSDLAG00005026832 (*ifi27l2a*) ($p$ = 0.927, r =0.448) in Cluster B. B) Boxplot showing the gene expression of miR-199-5p in animals with different VNN resistance QTL genotypes in Cluster B
